# Catestatin suppresses melanoma progression and drug resistance through multitargeted modulation of signaling pathways

**DOI:** 10.1101/2025.10.22.684062

**Authors:** Satadeepa Kal, Suborno Jati, Kechun Tang, Nicholas J.G. Webster, Angelo Corti, Sushil K. Mahata

## Abstract

**Background:** Despite advances in targeted and immune-based therapies, melanoma remains one of the most aggressive and treatment-resistant cancers. Resistance to small-molecule inhibitors and immune checkpoint blockade highlights the need for new mechanistically distinct interventions. Catestatin (CST), a Chromogranin A (CgA)– derived peptide with immunomodulatory and reparative properties, has been implicated in tissue protection, but its role in melanoma remains unknown.

**Methods:** CST expression was analyzed across melanoma stages and correlated with disease progression. Functional effects of CST were assessed in patient-derived and established melanoma cell lines, as well as in B16-F10 melanoma–bearing mice. RNA sequencing and pathway analyses were performed to delineate CST-regulated molecular networks. Vemurafenib-resistant A375 cells were used to examine CST’s effects on drug resistance mechanisms.

**Results:** CST expression declined with advancing tumor stage. CST treatment inhibited proliferation, migration, and invasion, while inducing apoptosis in melanoma cells but not in normal fibroblasts. *In vivo*, systemic CST administration significantly reduced tumor volume and mass. Transcriptomic profiling revealed coordinated downregulation of hypoxia-inducible, epithelial–mesenchymal transition (EMT), and collagen-remodeling pathways, alongside suppression of oxidative stress–adaptive signaling. In Vemurafenib-resistant A375 cells, CST restored apoptotic sensitivity and repressed multiple MAPK and PI3K–AKT–linked resistance genes.

**Conclusions:** CST acts as a mechanistically distinct peptide modulator that reprograms oncogenic signaling through inhibition of hypoxia, EMT, and survival pathways. These findings identify CST as a promising therapeutic prototype for mitigating melanoma progression and overcoming resistance to targeted therapy.

## INTRODUCTION

Melanoma is one of the most rapidly increasing cancers worldwide, with incidence rates rising faster than any other solid tumor over recent decades ^1,2^. Although it accounts for a small fraction of all skin cancer cases, melanoma is responsible for more than 80% of skin cancer–related deaths owing to its aggressive metastatic potential and resistance to therapy ^2^. The advent of targeted therapies—particularly BRAF and MEK inhibitors—and immune checkpoint blockade has substantially improved outcomes for patients with advanced disease ^3–5^. Small-molecule inhibitors such as Dabrafenib and Vemurafenib, which selectively target *BRAF^V600E* mutations, are now standard of care for a majority of melanoma patients ^5,6^, whereas those harboring *KIT* mutations benefit from Imatinib ^7^. Moreover, the introduction of immune checkpoint inhibitors has revolutionized melanoma management, extending median survival from approximately six months to nearly six years ^4,8^.

Despite these major advances, a significant proportion of patients either fail to respond or acquire resistance during treatment, resulting in relapse and mortality ^9,10^. These limitations highlight the urgent need for novel therapeutic approaches that engage mechanisms beyond those targeted by current modalities.

Chromogranin A (CgA), a prohormone abundantly expressed in neuroendocrine tissues ^11,12^, undergoes proteolytic processing to generate multiple biologically active peptides. Among them, Catestatin (CST; hCgA_352-372_) has emerged as a multifunctional regulator of cardiovascular, metabolic, and immune homeostasis ^13–17^. CST exerts broad protective effects, including anti-inflammatory ^18,19^, anti-hypertensive ^20–24^, and anti-diabetic ^18,25^ actions. Mechanistically, it inhibits catecholamine release ^26–28^, regulates endothelial proliferation and migration ^19,29,30^, and modulates macrophage and lymphocyte phenotypes ^18^, collectively supporting its role as a systemic homeostatic peptide with therapeutic potential.

CST is also expressed in human skin, where its levels increase following injury ^31^, and exerts functional activity in keratinocytes ^32^. CST-like immunoreactivity has been reported in rodent skin and sensory ganglia ^33^, suggesting a conserved cutaneous role. However, its significance in melanoma pathobiology has not been explored.

In the present study, we identify CST as a previously unrecognized regulator of melanoma progression in both human and mouse models. We demonstrate that CST expression decreases with advancing melanoma stage and that exogenous CST markedly suppresses melanoma cell viability, proliferation, and migration across patient-derived and established cell lines. In syngeneic B16-F10 tumor models, CST administration significantly reduces tumor growth and enhances apoptosis, confirming its antitumor efficacy *in vivo*. Transcriptomic analyses reveal that CST downregulates key pathways associated with extracellular-matrix organization, hypoxia response, and collagen metabolism—hallmarks of melanoma invasion and metastasis. Notably, CST suppresses several resistance-associated genes, including *FGFR3, PDGFRB, ID1/2/3, SREBF1,* and *MITF*, in both Vemurafenib-sensitive and Vemurafenib-resistant A375 cells, suggesting a role in overcoming therapeutic resistance.

Collectively, our findings establish CST as a novel melanoma-regulatory peptide with multifaceted antitumor activity. By integrating clinical observations, molecular analyses, and *in vivo* validation, this study positions CST as a promising therapeutic candidate capable of complementing existing targeted and immunotherapies. Given its pleiotropic regulatory properties and favorable safety profile, CST and its optimized analogs may inaugurate a new class of peptide-based therapeutics that modulate immune–metabolic networks to inhibit melanoma progression and resistance.

## RESULTS

### 1. Catestatin levels decline with advancing melanoma and exhibit anti-proliferative effects on patient-derived melanoma cells

Although CST has been detected in human skin ^31^ and keratinocytes ^32^ as well as in rodent skin and ganglia ^33^, its presence in melanoma patient samples had not been established. To address this, we analyzed CST protein expression across progressive stages of melanoma using tissue microarray slides (TissueArray.com). Immunohistochemical (IHC) analysis revealed a marked decline in CST expression with advancing melanoma stages, whereas normal skin and stage I melanoma tissues retained high CST levels (**Fig. 1A**). Quantitative analysis confirmed a significant reduction in CST staining intensity as melanoma progressed (**Fig. 1B**).

**Figure 1.**
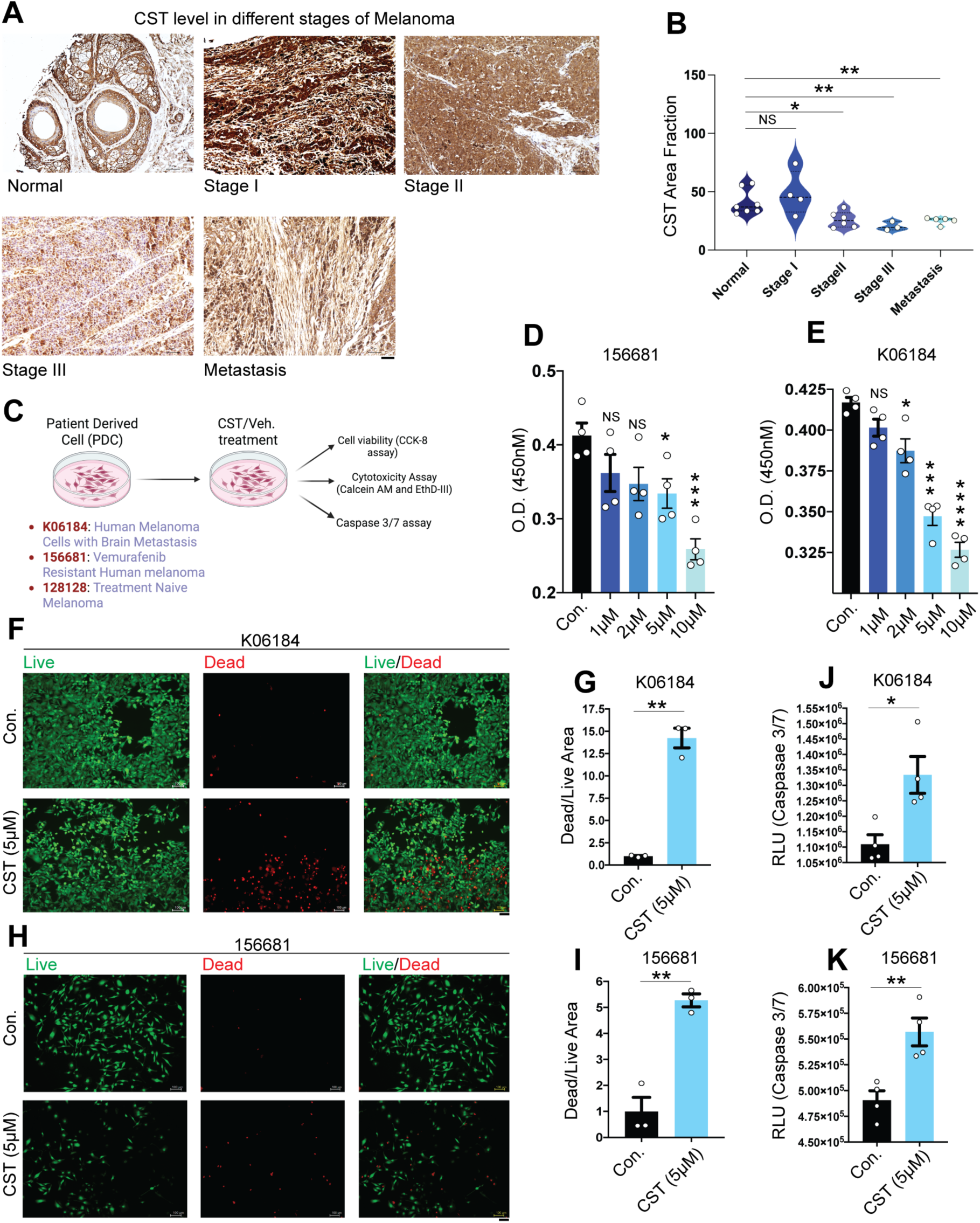
Decline in endogenous CST levels in human melanoma samples and melanoma cell proliferation inhibition upon extraneous CST treatment. **(A)** Representative immunohistochemistry images showing levels of Catestatin (CST) in different melanoma stages (normal, stage I, stage II, stage III, and metastasis) in human samples from TissueArray.com. **(B)** Quantification of the immunoreactive CST as area fraction as determined by Fiji3 software. (**C)** Schematic for experimental setup on melanoma patient derived cells (K06184, 156681, and 128128). (**D&E)** Cell viability assay performed on patient derived cell 156681 and K06184 for 120 hours in response to different concentrations of CST (1 µM, 2 µM, 5 µM, and 10 µM as compared to vehicle control (n=4). (**F&H)** Live and dead cell imaging of mammalian cells microscopy showing calcein AM (green) stained live cells and EthD-III (red) stained dead cells upon CST treatment versus control in K06184 and 156681 cells. Images were captured in Keyence microscope at a magnification of 10X. Scale bar for imaging is 100 µm. (**G&I)** Dead/Live area fraction measured upon CST treatment (5 µM) in K06184 and 56681 cells (n=3). (J&K) Caspase 3/7 assay done in CST-treated and untreated K06184 and 156681 cells. (n=4). Data were presented as Mean ± SEM. Statistical analyses were done using one-way ANOVA (D&E) and Welch’s t-test (B,G,J,I,K). \**p* ≤ 0.05, \*\**p* ≤ 0.01, \*\*\**p* ≤ 0.001, \*\*\*\**p* ≤ 0.0001.

To determine whether CST administration affects melanoma cell viability, we used patient-derived melanoma cell lines K06184, 156681, 128128 (characteristics mentioned in **Supplementary File 1**) from the NCI-PDMR repository and normal human dermal fibroblasts (CCD1076). A schematic of the experimental plan with the patient derived cell lines are mentioned (**Fig. 1C**). CST treatment 1 µM, 2 µM, 5 µM, 10 µM) for 120 hours caused dose-dependent reduction in melanoma cell viability (**Fig. 1D& E; Supplementary Fig. S1B**), whereas normal fibroblasts remained unaffected even at the highest concentration (**Supplementary Fig. S1A**). Fluorescence-based viability/cytotoxicity assays showed a pronounced increase in dead cells (EthD-III, red) relative to live cells (Calcein-AM, green) following CST treatment (**Fig. 1F-H**; **Supplementary Fig. S1C**). Quantification revealed a significantly elevated dead-to-live cell ratio (**Fig. 1G-I, Supplementary Fig. S1D**). Corroboration of these results by increased caspase-3/7 activity not only confirm induction of apoptosis (**Fig. 1J-K, Supplementary Fig. S1E**), but hint at a possible role of CST in curbing melanoma progression.

### 2. Catestatin reduces viability, proliferation, and migration in melanoma cell lines

To further our preliminary observations regarding the role of CST in curbing melanoma, we performed several experiments in established mouse and human melanoma cell lines and their normal counterparts as shown in schematic (**Fig. 2A**). We performed cell viability assay in mouse melanoma cell line B16-F10 and human melanoma cell lines A375, SKMEL28 with increasing concentration of CST (0.5 µM, 1 µM, 2 µM, and 5 µM) for 72 hours. While we recorded a marked decrease in cell viability of the melanoma cell lines (**Fig. 2C, E&F**), there was no significant reduction in cell viability of normal mouse fibroblast or human skin fibroblast cell lines (**Fig. 2B&D**), thus indicating that CST preferentially kills the cancer cells without affecting the normal ones. Based on the maximal effects, a concentration of 2 µM was used for subsequent assays.

**Figure. 2.**
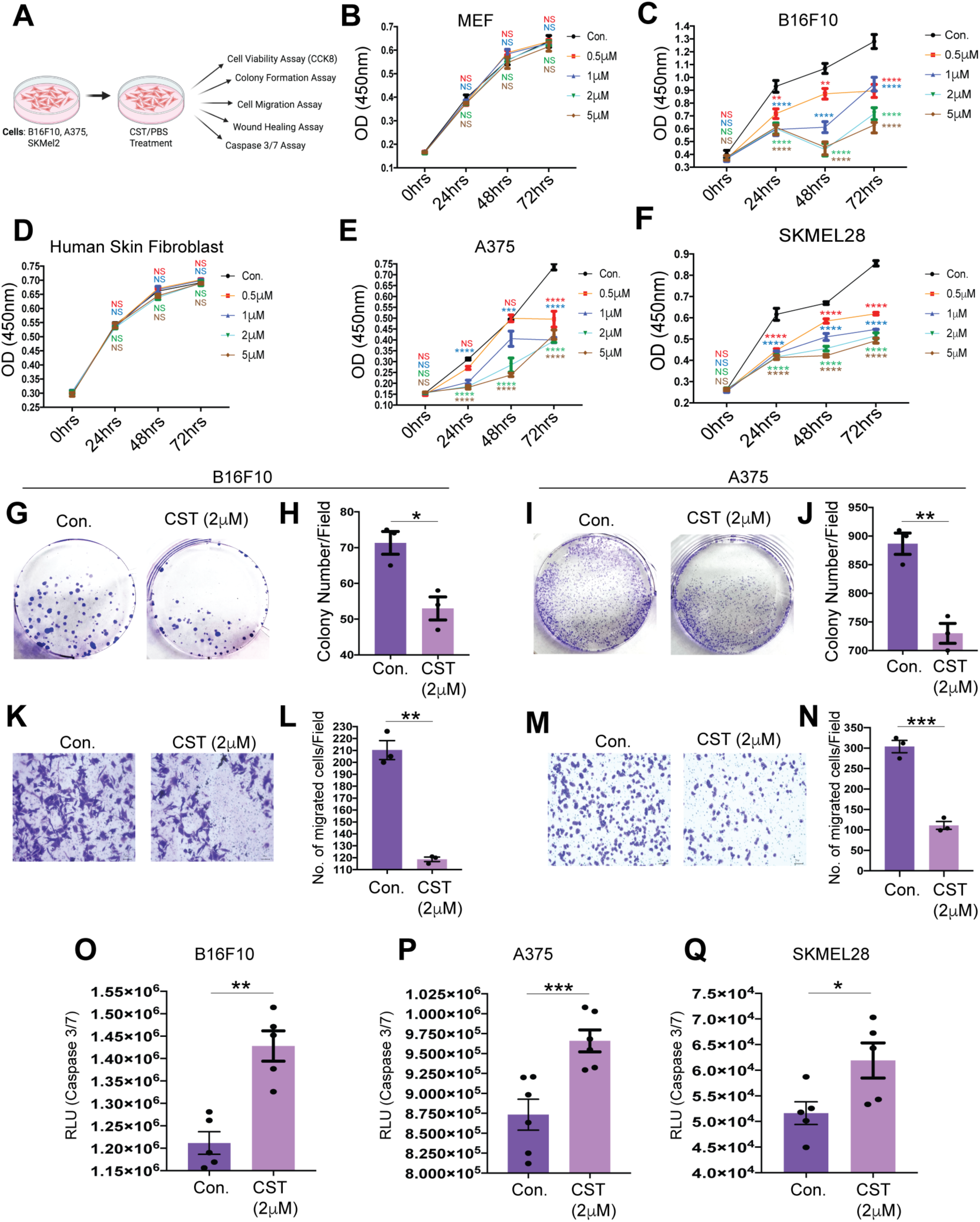
Melanoma progression inhibition upon CST treatment in mouse and human melanoma cell lines. **(A)** Schematic representation of experimental setup on B16F10, A375 and SKMEL28 cell line in control and CST-treated condition. (**B&C)** Cell viability of mouse embryonic fibroblast (MEF) and mouse melanoma cell line B16F10 by CCK-8 assay using different concentrations of CST (0.5 µM, 1 µM, 2 µM, and 5 µM) versus vehicle control for 0, 24,48 and 72 hours. (**D-F)** Cell viability by CCK-8 assay on normal human skin fibroblast and melanoma cell lines A375,SKMEL28 in response to different concentrations of CST (0.5 µM, 1 µM, 2 µM, and 5 µM) for 0, 24, 48 and 72 hours. For all cck-8 assay (n=4). (**G&H)** Colony formation assay and its quantitative analysis in B16F10 mouse melanoma cells. **(I&J**) Colony formation and its quantitative analysis in A375 human melanoma cell (n=3). (**K&L**) Transwell migration assay in CST versus vehicle-treated condition and its quantitative analysis in B16F10 cells. (**M-N)** Transwell migration assay in CST versus vehicle-treated condition along with quantitative analysis in A375 human melanoma cells (n=3). These bright field images were captured in Keyence microscope at 10X magnification. Scale bar: 100 µm. (**O-Q)** Caspase3/7 Assay in CST versus vehicle-treated condition in B16F10, A375 and SKMEL28 cells (n=5). Data were presented as Mean ± SEM. Two-way ANOVA followed by Dunnett’s multiple comparison test was used to analyze cell viability assay data. Welch’s t-test was used to analyze colony formation, transwell migration and Caspase3/7 assay data. \**p* ≤ 0.05, \*\**p* ≤ 0.01, \*\*\**p* ≤ 0.001, and \*\*\*\**p* ≤ 0.0001.

Phase-contrast microscopy revealed increased cell death in A375 cells treated with CST for 24 hours (**Supplementary Fig. S2A**). Colony-formation assays showed that CST markedly suppressed clonogenic potential in both human and mouse melanoma cells (**Fig. 2G–J**; **Supplementary Fig. S2B&C**). Because cell migration is a hallmark of metastatic potential, we evaluated migration using transwell and wound-healing assays. CST significantly impaired migration in all melanoma lines compared with controls (**Fig. 2K–N**; **Supplementary Fig. S2D–G**). Increased caspase-3/7 activity in CST-treated melanoma cells (**Fig. 2O–Q**) further confirmed apoptotic induction. Collectively, CST inhibits melanoma cell survival, proliferation, and metastatic potential.

### 3. Catestatin reduces melanoma tumor burden *in vivo*

To assess CST efficacy *in vivo*, C57BL/6 mice were subcutaneously injected with 5 × 10⁴ B16-F10 melanoma cells and treated intraperitoneally with CST (10 mg/kg, three times per week) (**Fig. 3A**). Our *in vitro* plasma stability studies revealed the following remaining percentages of CST during the 24 hours of incubation period: 58.9% after 0.125 hour; 20.2% after 8 hours; 0% after 24 hours (**Supplementary Fig. S3A**). We found the following in vivo plasma pharmacokinetic values: *C*_max_ (maximum concentration): 148 ng/ml; *T*_max_ (time to reach Cmax): 0.25 hour; *t*_1/2_ (half-life): 1.38 hour (**Supplementary Fig. S3B**). CST treatment significantly suppressed tumor growth compared with vehicle controls (**Fig. 3B**). No major changes in body weight or liver histology were observed (**Supplementary Fig. S3C&D**), indicating systemic tolerability. The tumor volume kinetics revealed marked decrease upon CST treatment and tumor weight was also lower in CST-treated mice (**Fig. 3C&D**). IHC staining revealed reduced Ki-67 expression, indicating diminished proliferation (**Fig. 3E&F**). Hematoxylin and eosin staining showed reduced cellular density (**Fig. 3E**), and TUNEL assay demonstrated increased apoptotic nuclei in CST-treated tumors (**Fig. 3G&H**). Western blot analysis confirmed elevated cleaved caspase-3 levels (**Fig. 3I&J**). Thus, CST treatment significantly curtails melanoma growth *in vivo* without observable toxicity.

**Figure 3.**
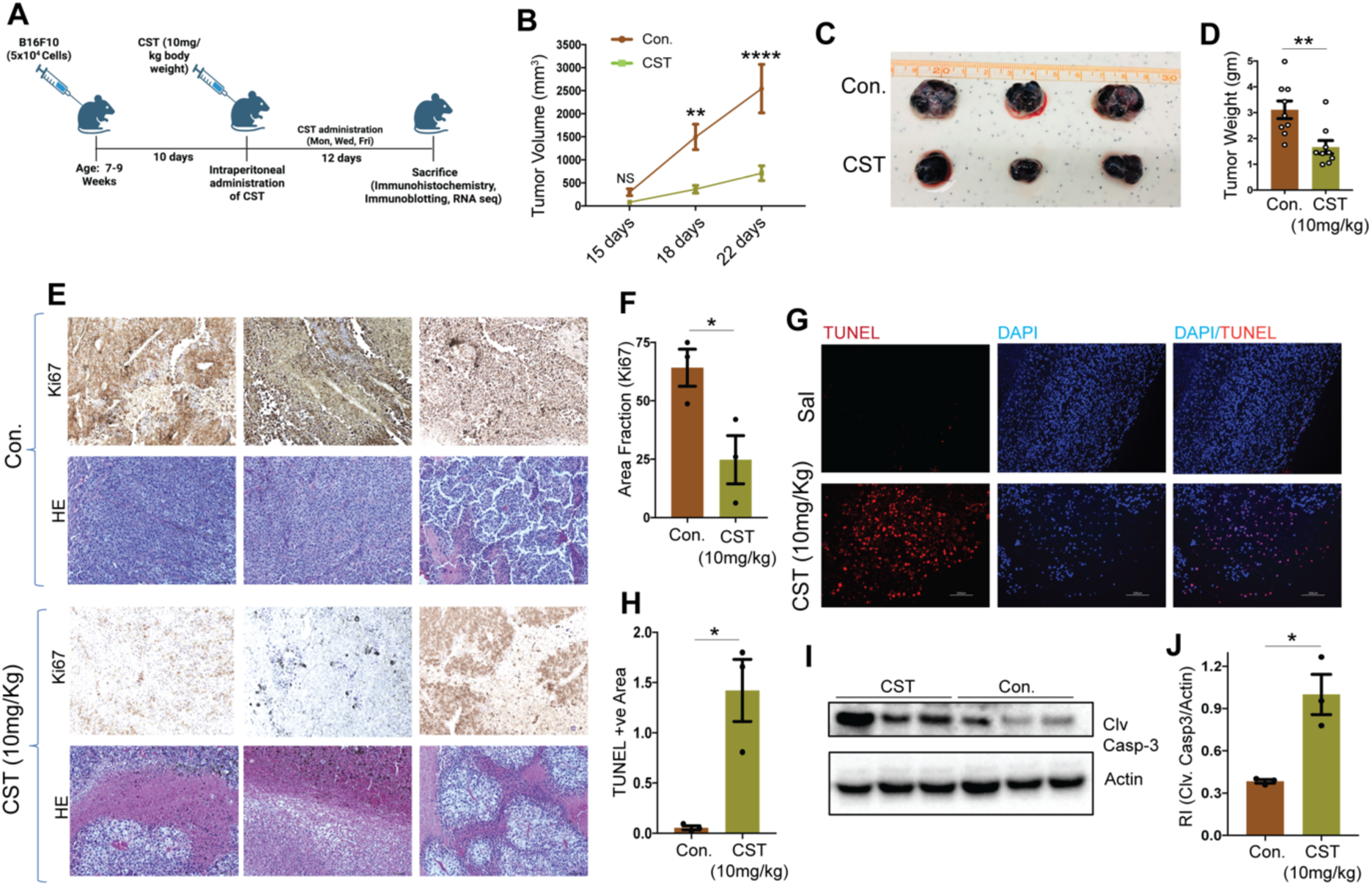
***In vivo* melanoma tumor is reduced upon CST treatment** (**A**) Schematic showing injection of 5*10^^4^ B16F10 cells to 7-9 weeks old C57BL/6 mice and administration of CST (10 mg/kg) thrice a week till sacrifice. (**B)** Tumor growth kinetics in B16F10 derived tumors treated with CST (10mg/kg) and vehicular control (n=9). (**C)** Post-harvesting images of Control and CST-treated tumors. (**D)** Bar graph showing changes in tumor weight in response to control or treatment with CST (n=9). (**E)** Immunohistochemistry images showing levels of proliferation marker Ki-67 in Control and CST treated mice tumors. Hematoxylin and eosin staining reveal the tissue structures of the tumors. Images were captured in brightfield in Keyence microscope at a magnification of 20X. Scale bar: 50 µm. (**F)** Bar graph showing area fraction of Ki-67 staining in control and CST-treated tumors (n=3). (**G)** TUNEL assay performed on control and CST treated tumor sections. The TUNEL positive cells stain red and the nuclei stain blue. Images were captured in a Keyence microscope at magnification of 10X. Scale bar: 100 µm. (**H**) The TUNEL-positive area of different tumors are represented in the graph (n=3). (**I)** Representative western blot of cleaved caspase3 as compared to actin levels in mice after treatments with control or CST treated tumors. (**J**) Bar graph showing densitometric analysis of I(n=3). Data were presented as Mean ± SEM and analyzed by 2-way ANOVA followed by Tukey’s multiple comparison test (tumor growth kinetics) or by Welch’s t-test. \**p* ≤ 0.05, \*\**p* ≤ 0.01, and \*\*\*\**p* ≤ 0.001.

### 4. Molecular mechanisms underlying CST-mediated melanoma regression

To elucidate the mechanisms of CST-induced tumor suppression, bulk RNA sequencing was performed on control and CST-treated B16-F10 tumors (n = 4 per group). Differential expression analysis revealed widespread transcriptional modulation (**Fig. 4A**) with 184 significantly upregulated and 157 significantly downregulated genes upon CST treatment *in vivo*. Gene-ontology enrichment upregulated (**Supplementary Fig. S4A&B**) and downregulated transcripts were mapped and when emphasized on the downregulated pathways they were linked to *extracellular matrix (ECM) organization*, *collagen metabolism*, *response to hypoxia*, and *epithelial-to-mesenchymal transition (EMT)* (**Fig. 4B&C**)—processes that facilitate melanoma progression ^34–36^.

**Figure. 4.**
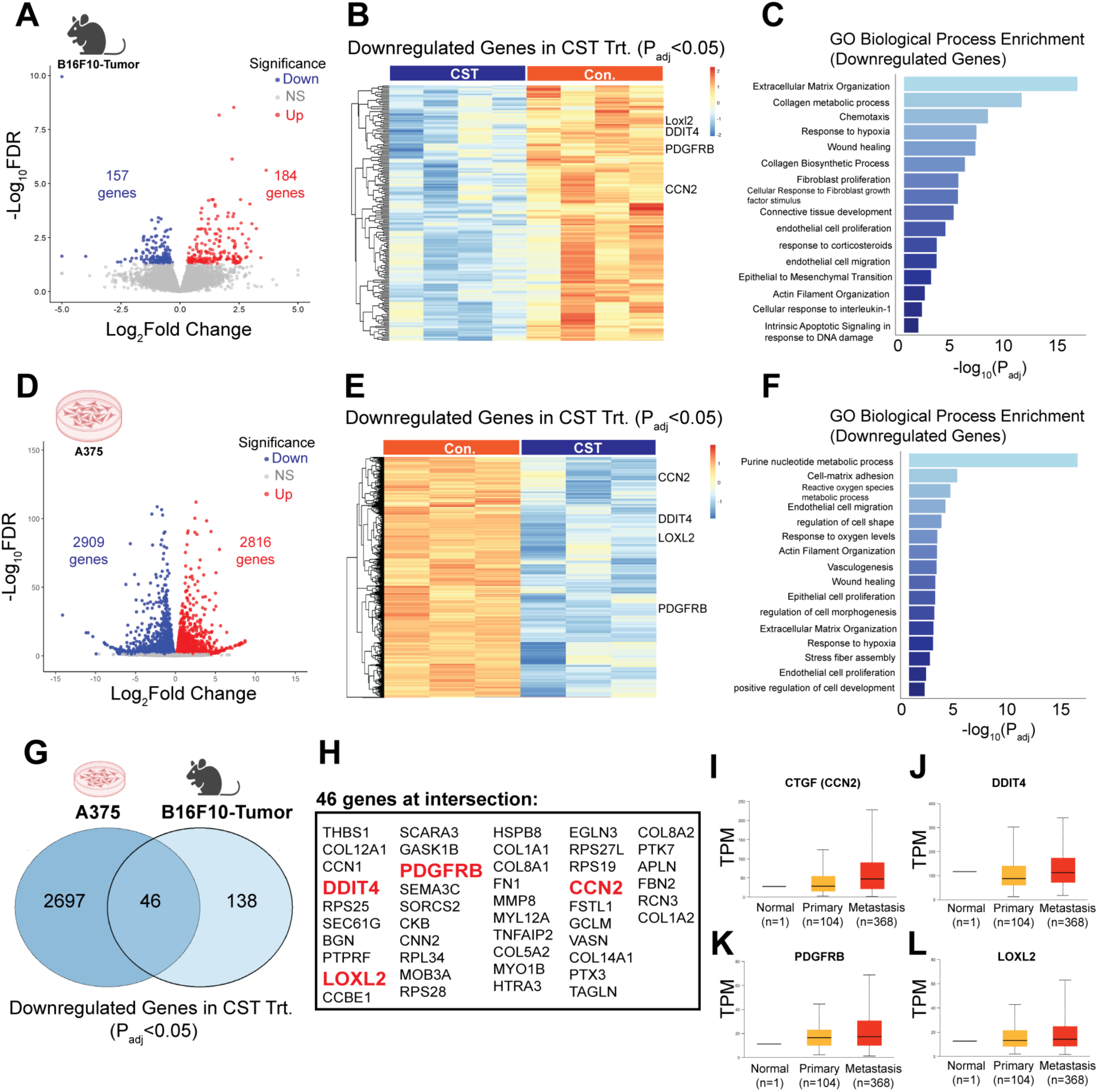
Molecular mechanisms driving CST function in mouse tumor and human melanoma cell line. (**A**). Volcano plot of differential regulation of genes upon CST treatment. (**B)** Heat map showing all downregulated genes with padj <0.05 with emphasis to *CCN2*, *LOXL2*, *DDIT4*, *PDGFRB*, and FN1 genes (n=4, each group) (**C**) Gene ontology (GO) analysis of enriched downregulated pathways emphasizing on the previously mentioned genes. (**D)** Volcano plot of differentially expressed genes upon CST treatment in A375 human melanoma cell line. (**E)** Heatmap of downregulated genes with padj <0.05 (n=3, each group) (**F)** GO analysis of enriched downregulated pathways upon CST treatment in A375 cells. (**G)** Venn Diagram showing common downregulated genes in B16F10 derived mouse tumor and A375 cell line. (**H)** List of the 46 common genes in the Venn Diagram with emphasis on *CCN2*, *LOXL2*, *DDIT4*, *PDGFRB*, and *FN1*. (**I-L)** Expression levels of *CCN2*, *DDIT4*, *PDGRB* and *LOXL2* from TCGA database in primary melanoma vs metastasis as analysed in UALCAN database.

Similarly, transcriptomic analyses of CST-treated A375 cells showed 2816 upregulated genes and 2909 downregulated genes (**Fig. 4D**) The upregulated (**Supplementary Fig. S4C&D)** and downregulated genes were mapped and the downregulated pathways related to *cell-substrate adhesion*, *reactive oxygen species metabolism*, and *hypoxia response* (**Fig. 4E-F**) were also observed in human melanoma A375 cell line. Comparative analysis of the mouse and human cell line datasets revealed 46 commonly downregulated genes (**Fig. 4G&H**), including *CCN2*, *LOXL2*, *DDIT4*, *PDGFRB* and *FN1*, all previously implicated in melanoma aggressiveness ^37–40^. Consistent with these findings, TCGA-SKCM analyses via UALCAN confirmed elevated expression of these genes in advanced melanoma stages (**Fig. 4I–L**). Together, these results suggest that CST suppresses melanoma by downregulating ECM remodeling, hypoxia, and EMT-associated signaling programs.

### 5. CST suppresses Vemurafenib-resistant melanoma

Resistance to BRAF inhibitors such as Vemurafenib remains a major clinical challenge. Notably, the patient-derived melanoma line 156681 (detailed in **Supplementary File 1**), isolated from a Vemurafenib-treated non-responder, displayed enhanced CST sensitivity with decreased viability and increased cell death as observed previously (**Fig. 1**). This observation led us to check further the potential of CST as an anti-cancerous agent in Vemurafenib resistant cells. To validate this observation, we generated a Vemurafenib-resistant A375 cell line. The IC₅₀ for resistant cells (1432 nM) was >6-fold higher than that of parental cells (228.9 nM) (**Fig. 5A&B**). CST treatment (0.5 µM, 1 µM, and 2 µM) markedly reduced viability of resistant A375 cells (**Fig. 5C**) and induced extensive cytopathic changes (**Fig. 5D**). Transwell migration assays confirmed diminished migratory capacity following CST exposure (**Fig. 5E**). To further into the molecular mechanism of these observed effects, transcriptomics was performed in control and CST-treated resistant A375 cells.

**Figure 5.**
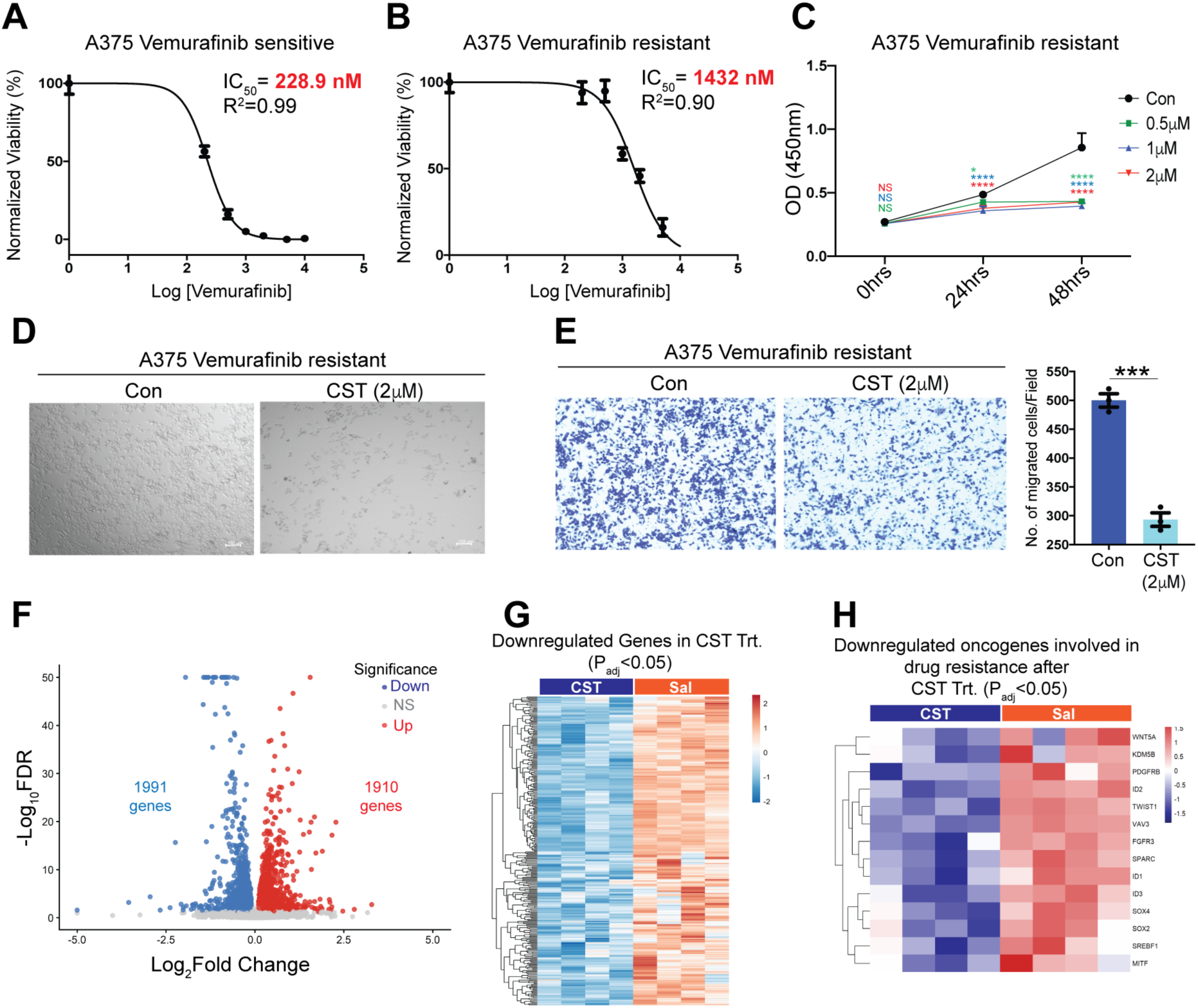
CST kills Vemurafenib resistant A375 melanoma cells by diminishing the levels of resistance associated genes. **(A)** IC_50_ of Vemurafenib in Vemurafenib-sensitive A375 cell line. (**B)** IC_50_ of Vemurafenib in Vemurafenib-resistant A375 cell line. (**C)** Dose-dependent (0.5 µM, 1 µM, 2 µM, and 5 µM) effects of CST on cell viability of Vemurafenib-resistant A375 cells after 0,24 and 48 hours of treatment (n=6). (**D)** Phase contrast image of control versus CST-treated Vemurafenib resistant A375. (**E)** Bar graph showing quantitative analysis of Trans well migration assay in Vemurafenib-resistant A375 cells (n=3). (**F**) Volcano plot of differentially expressed genes upon CST treatment in A375 resistant cell (n=4, each group) **(G)** Heatmap showing downregulated genes upon CST treatment with padj <0.05. (**H)** Heatmap of downregulated oncogenes with known link to resistance towards standard melanoma treatment including *WNT5A*, *KDM5B*, *PDGFRB*, *ID1*, *ID2*, *ID3*, *TWIST1*, *VAV3*, *FGFR3*, *SPARC*, *SOX2*, *SOX4*, *SREBP1*, and *MITF*. Cell viability assay graphs were analysed using 2-way ANOVA followed by Dunnett’s multiple comparison test and Welch’s t test for analysis of the transwell migration data. \**p* ≤ 0.05, \*\**p* ≤ 0.01, and \*\*\**p* ≤ 0.001.

Transcriptomic analysis of these resistant cells treated with CST revealed extensive transcriptional reprogramming (**Fig. 5F&G, Supplementary Fig. S5A&B**) with 1990 upregulated genes and 1991 downregulated genes, including downregulation of *WNT5A* ^41^*, PDGFRB* ^42^*, ID1–3* ^43–45^*, FGFR3* ^46^*, SREBF1* ^47^*, VAV3*^48^*, KDM5B* ^49–51^*, TWIST1* ^52^*, SOX2* ^53^*, SOX4* ^54^*, MITF* ^55^, and *SPARC* ^56^ (**Fig. 5H**)—genes associated with resistance and metastasis. The suppression of these genes that are actively involved in conferring resistance to known treatment regimens underscores the therapeutic potential of CST in overcoming treatment-resistant melanoma.

## DISCUSSION

Melanoma incidence continues to rise worldwide, with an estimated 331,647 new cases and 58,645 deaths reported in 2022 (GLOBOCAN 2022) ^57^. In the United States alone, over 104,000 new invasive melanoma cases are projected for 2025 (https://www.curemelanoma.org/blog/over-104-000-americans-estimated-to-be-diagnosed-with-invasive-melanoma-in-2025). Despite major advances in targeted and immune-based therapies, recurrence and therapy resistance remain formidable clinical barriers ^58^. This persistent challenge underscores the need for novel, mechanistically distinct therapeutic strategies capable of curbing melanoma progression and overcoming resistance to current regimens.

In this study, we introduce CST as a promising therapeutic candidate against melanoma. Peptide-based therapeutics are underexplored in this malignancy ^59–62^, yet they offer inherent advantages, including biocompatibility, low immunogenicity, and ease of modification. CST, known for its anti-inflammatory ^18,24^ and cardio-metabolic benefits ^13^, emerged as a rational candidate owing to its regulatory roles in inflammation, oxidative stress, and metabolic homeostasis—processes intimately linked to oncopathobiology.

Our analyses reveal that CST expression declines progressively with melanoma stage, suggesting its loss may facilitate tumor advancement. Restoration of CST in patient-derived melanoma cells elicit marked cytotoxicity and apoptosis in lines with distinct molecular features—K06184 (wild-type BRAF, brain metastasis) and 156681 (BRAF^V600E, Vemurafenib-non-responder) and 128128 (treatment naïve, metastatic melanoma)—highlighting its broad anti-cancer efficacy across genotypes.

Mechanistically, CST exerted potent anti-proliferative and anti-migratory effects. In both mouse (B16F10) and human (A375, SKMEL28) melanoma cell lines, CST treatment reduced cell viability, colony formation, and transwell migration, while sparing normal fibroblasts. *In vivo*, CST administration markedly suppressed tumor growth in B16F10 melanoma-bearing C57BL/6 mice, with concomitant reductions in Ki-67 and elevations in TUNEL and cleaved-caspase-3 signals. The absence of changes in body weight or hepatic histology confirmed that CST is well tolerated and systemically non-toxic.

To define CST-driven molecular alterations, transcriptomic analyses identified widespread transcriptional reprogramming. Downregulated gene clusters were enriched in pathways controlling *extracellular-matrix organization*, *collagen metabolism*, *hypoxia adaptation*, and *epithelial-to-mesenchymal transition (EMT)*—all central to melanoma invasion and metastasis. CST significantly suppressed *CCN2, LOXL2, DDIT4,* and *FN1*, key drivers of ECM remodeling and angiogenesis, indicating that CST disrupts the structural and metabolic axes sustaining tumor progression.

Given the high prevalence of therapeutic resistance, particularly to BRAF/MEK inhibitors, we further evaluated any potential role of CST in Vemurafenib-resistant A375 cells. CST markedly reduced cell viability and migration in these resistant cells, accompanied by downregulation of multiple resistance-associated genes—including *FGFR3* ^46,63^, *ID1/2/3* ^43,64^, *PDGFRB*, *SREBF1* ^47,65^, and *MITF* ^66^. These targets are central mediators of adaptive survival and dedifferentiation in BRAF^V600E melanoma, suggesting that CST not only restrains tumor growth but may re-sensitize resistant cells to existing therapies.

Collectively, our findings delineate CST as a multi-targeted anti-melanoma peptide that impairs proliferation, invasion, and resistance mechanisms while maintaining a favorable safety profile. Although CST has been variably reported to promote angiogenesis in other contexts ^30,67^, our *in-vivo* and transcriptomic data consistently support an anti-angiogenic and anti-proliferative role in melanoma.

Future directions include optimizing CST pharmacodynamics through peptidomimetic modifications to enhance stability, half-life, and tumor bioavailability. Combination therapy paradigms integrating CST with BRAF/MEK inhibitors or immune checkpoint blockade warrant exploration to achieve synergistic efficacy. Moreover, since Chromogranin A yields several bioactive fragments—Pancreastatin ^68^, Vasostatin I/II ^69^, WE14 ^70^, and Serpinin ^71^—systematic studies of their processing and interactions in melanoma may reveal complementary or cooperative functions.

In conclusion, this study provides compelling preclinical evidence that Catestatin represents a novel, safe, and effective therapeutic modality for both treatment-naïve and drug-resistant melanoma. By simultaneously targeting tumor growth, metastatic potential, and resistance pathways, CST holds significant translational promise as the foundation for a new class of peptide-based anti-melanoma therapies.

## Acknowledgments

This work was supported by NIH grants AG080246, AG078635, and AG091126 to SKM as well as VA RR&D SPiRE grant RX004398 to SKM. This work was also supported by a VA Merit Review Award (I01BX004848) and Senior Research Career Scientist Award (IBX005224) to NW.

## Contribution to work

Conceptualization: SKM, SK Methodology: SK, SJ, KT, SKM, AC Investigation: SK

Visualization: SK, SJ, SKM, NW Funding acquisition: SKM, NW Project administration: SKM, SK, SJ Supervision: SK, SJ, SKM

Writing – original draft: SK, SKM

## MATERIALS AND METHODS

### Cell Culture

Mouse embryonic fibroblast (MEF), human skin fibroblast (CCD1076), human melanoma cell lines (A375, SK-MEL-28), and mouse melanoma cell line (B16-F10) were obtained from the American Type Culture Collection (ATCC). Cells were maintained in DMEM supplemented with 10% fetal bovine serum (FBS), 100 U/mL penicillin, and 100 µg/mL streptomycin in a humidified incubator at 37°C with 5% CO₂. Patient-derived melanoma cell lines (K06184, 156681, and 128128) were procured from the NCI Patient-Derived Models Repository (PDMR) and cultured following PDMR standard operating procedures in DMEM/F-12 containing 5% FBS, 0.4 µg/mL hydrocortisone, 0.01 µg/mL epidermal growth factor, 24 µg/mL adenine, 100 µg/mL penicillin-streptomycin, 2 mM L-glutamine, and 10 nM Y-27632 dihydrochloride on Matrigel-coated plates.

### Human Tissue Samples

Commercially available human melanoma tissue microarrays (catalog nos. Me482A and Me551) were purchased from TissueArray.com for immunohistochemical analyses.

### Peptide Treatment

Catestatin (**CST**; sequence SSMKLSFRARAYGFRGPGPQL) was synthesized by GenScript and dissolved in 0.9% saline. Cells were treated with CST at concentrations of 0.5–10 µM as indicated. For *in vivo* studies, CST was administered intraperitoneally at 10 mg/kg body weight, three times per week (Monday, Wednesday, Friday).

### Bioinformatic Analyses

Expression of CCN2, LOXL2, DDIT4, PDGFRB **and** FN1 in primary and metastatic skin cutaneous melanoma samples was analyzed using the UALCAN database (http://ualcan.path.uab.edu) ^72^.

### Caspase Assay

Cells (1 × 10⁴ per well) were seeded in 96-well white plates and treated with or without CST for 48 h (A375, SK-MEL-28, B16-F10) or 120 h (patient-derived lines K06184, 128128, 156681). Caspase-3/7 activity was measured using the Caspase-Glo® 3/7 Assay System (Promega) following the manufacturer’s protocol. Luminescence was recorded using a Varioskan LUX microplate reader (Thermo Fisher Scientific).

### Live/Dead Viability Assay

Cells were treated with CST or vehicle for 120 h, washed with PBS, and incubated with 2 µM calcein AM and 4 µM ethidium homodimer III (Biotium Viability/Cytotoxicity Assay Kit) for 30 min at room temperature in the dark. After washing, cells were imaged under FITC and Texas Red® channels using a Keyence fluorescence microscope (20×).

### Cell Viability Assay

Cells (1 × 10⁴ per well) were plated in 96-well plates and treated with CST (0.5–10 µM) for 72–120 h. Viability was determined using the CCK-8 reagent (APExBIO). Absorbance was measured at 450 nm using a Varioskan LUX reader.

### Colony Formation Assay

A375, SK-MEL-28, and B16-F10 cells (1 × 10³ per well) were treated with or without CST according to the experimental design and cultured for 15 days. Colonies were fixed in methanol for 20 min, stained with 0.1% crystal violet, and photographed.

### Transwell Migration Assay

Cells pretreated with CST or vehicle for 48 h were seeded in serum-free medium in transwell inserts (8 µm pore size). The lower chamber contained complete medium. After 24 h, non-migrated cells were removed, membranes were fixed, stained with crystal violet, and imaged using a Keyence brightfield microscope (20×) as described previously ^73^.

### Wound Healing Assay

A375 cells at 40–50% confluency was scratched with a sterile pipette tip, followed by CST or control treatment. Images were captured at 0 and 24 h using Keyence brightfield microscopy (20×).

### RNA Sequencing

Total RNA from mice tumor and A375 cell line was isolated using RNeasy miniprep Kit (Qiagen). RNA was quantified by Nanodrop spectrophotometer and its integrity was evaluated by Tapestation (Agilent). Complementary DNA library preparation was performed using 400ng of RNA using mRNA HyperPrep Kit (KAPA) according to manufacturer’s protocol with Unique Dual-Indexed adapters (KAPA). The cDNA library was amplified and assessed by Qubit2.0 (Thermo Fisher Scientific). The libraries were then pooled and analysis was done in NovaSeq X Plus 10B (Illumina) in UCSD IGM core as described previously ^74^.

### Generation of Vemurafenib-Resistant Cells

Vemurafenib-resistant A375 cells were generated following established protocol. Parental cells were treated with 100 nM Vemurafenib for 2 weeks, followed by stepwise doubling of drug concentration every 2 weeks up to 3.2 µM. Surviving cells in maintenance culture containing 2 µM drug was used for downstream studies.

### Mouse Tumor Model

Age-matched male and female *C57BL/6* mice (7–9 weeks, n = 9) were subcutaneously injected with 5 × 10⁴ B16-F10 cells in 100 µL PBS. Ten days post-injection, CST (10 mg/kg) was administered intraperitoneally three times weekly. Tumor dimensions were measured with calipers, and volume was calculated using the formula V = (length × width²)/2. Mice were euthanized 22 days post-inoculation. All procedures were approved by the IACUC of UCSD and the VA San Diego Healthcare System and conformed to NIH guidelines.

### Immunohistochemistry and Histology

Tumors were fixed, paraffin-embedded, and sectioned (5 µm). Hematoxylin and eosin staining was performed at the La Jolla Institute for Immunology. Immunohistochemistry was carried out using the Super Sensitive™ M Polymer-HRP Kit (Biogenex). After deparaffinization, antigen retrieval was performed using Histo VT (Nacalai) at 90°C for 20 min. Sections were incubated overnight at 4°C with anti–Ki-67 (Abcam) or anti-CST (gift from Angelo Corti) at 1:100 dilution. Detection was performed with DAB and counterstained with hematoxylin. Slides were dehydrated and mounted in DPX. Images were captured using Keyence brightfield microscopy (20×).

### TUNEL Assay

Apoptotic nuclei were detected in tumor sections using the CF®594 TUNEL Assay Kit (Biotium). After deparaffinization and rehydration, sections were permeabilized with 20 µg/mL Proteinase K for 1 h at 37°C, incubated with TUNEL reaction mix for 2 h, counterstained with Hoechst, and mounted in antifade medium. Fluorescence images were obtained using a Keyence microscope.

### Phase-Contrast Microscopy

Morphological alterations in CST-treated A375 and Vemurafenib-resistant A375 cells were assessed using Keyence phase-contrast microscopy.

### Western Blotting

Tumor lysates were prepared in RIPA buffer containing protease inhibitors (Thermo Fisher). Equal protein amounts (10 µg) were resolved by SDS-PAGE, transferred to PVDF membranes, and incubated with antibodies against cleaved caspase-3 (Proteintech) and β-actin (Cell Signaling Technology). HRP-conjugated secondary antibodies (Cell Signaling Technology) were used, and signals were detected by chemiluminescence. Densitometric analyses were performed using GelQuant.NET (BiochemLabSolutions.com).

***In vitro* plasma stability and *in vivo* plasma pharmacokinetics**

Plasma stability was determined in 5 µg CST/ml spiked mouse (C57BL/6) plasma (50 µl) and taking aliquots at specific times over a 24-hour period. Each aliquot (50 µl) was mixed with 150 µl methanol, vortexed, spun, removed the supernate and reconstituted the pellet with 150 µl 10% formic acid in water. The sample was vortexed and spun and a 100 µl aliquot was taken, diluted 1:1 with 10 mM ammonium bicarbonate, and analyzed by LC-MS/MS.

For *in vivo* plasma pharmacokinetics, male C57BL/6 mouse received a 5mg/kg intraperitoneal dose of CST. Blood samples (100 µl) were obtained at 0, 0.25, 0.5, 1, 2, 4, 8, and 24 hours via tail vein bleeds. Plasma concentrations of CST was determined by LC-MS/MS methods as described above.

### Statistical Analyses

Data were analyzed using GraphPad Prism 10. Two-tailed Welch’s *t*-test was used for single-variable comparisons, and two-way ANOVA for multivariate datasets (e.g., concentration and time). Statistical significance was defined as \**p* ≤ 0.05*, **p ≤ 0.01**, \*\*\***p ≤ 0.001*, \*\*\**p ≤ 0.0001* and non-significant as n.s.

## REFERENCES

1. Whiteman DC, Green AC, Olsen CM. The Growing Burden of Invasive Melanoma: Projections of Incidence Rates and Numbers of New Cases in Six Susceptible Populations through 2031. J Invest Dermatol. 2016;136(6):1161–1171.

2. Siegel RL, Miller KD, Wagle NS, Jemal A. Cancer statistics, 2023. CA Cancer J Clin. 2023;73(1):17–48.

3. Robert C, Schachter J, Long GV, et al. Pembrolizumab versus Ipilimumab in Advanced Melanoma. N Engl J Med. 2015;372(26):2521–2532.

4. Hodi FS, O’Day SJ, McDermott DF, et al. Improved survival with ipilimumab in patients with metastatic melanoma. N Engl J Med. 2010;363(8):711–723.

5. Chapman PB, Hauschild A, Robert C, et al. Improved survival with vemurafenib in melanoma with BRAF V600E mutation. N Engl J Med. 2011;364(26):2507–2516.

6. Hauschild A, Grob JJ, Demidov LV, et al. Dabrafenib in BRAF-mutated metastatic melanoma: a multicentre, open-label, phase 3 randomised controlled trial. Lancet. 2012;380(9839):358-365.

7. Hodi FS, Corless CL, Giobbie-Hurder A, et al. Imatinib for melanomas harboring mutationally activated or amplified KIT arising on mucosal, acral, and chronically sun-damaged skin. J Clin Oncol. 2013;31(26):3182–3190.

8. Larkin J, Chiarion-Sileni V, Gonzalez R, et al. Five-Year Survival with Combined Nivolumab and Ipilimumab in Advanced Melanoma. N Engl J Med. 2019;381(16):1535–1546.

9. Kakadia S, Yarlagadda N, Awad R, et al. Mechanisms of resistance to BRAF and MEK inhibitors and clinical update of US Food and Drug Administration-approved targeted therapy in advanced melanoma. Onco Targets Ther. 2018;11:7095–7107.

10. Sharma P, Hu-Lieskovan S, Wargo JA, Ribas A. Primary, Adaptive, and Acquired Resistance to Cancer Immunotherapy. Cell. 2017;168(4):707–723.

11. Winkler H, Fischer-Colbrie R. The chromogranins A and B: the first 25 years and future perspectives. Neuroscience. 1992;49(3):497–528.

12. Bartolomucci A, Possenti R, Mahata SK, Fischer-Colbrie R, Loh YP, Salton SR. The extended granin family: structure, function, and biomedical implications. Endocr Rev. 2011;32(6):755–797.

13. Mahata SK, Kiranmayi M, Mahapatra NR. Catestatin: A Master Regulator of Cardiovascular Functions. Curr Med Chem. 2018;25(11):1352–1374.

14. Bandyopadhyay GK, Mahata SK. Chromogranin A Regulation of Obesity and Peripheral Insulin Sensitivity. Front Endocrinol (Lausanne). 2017;8:20.

15. Muntjewerff EM, Dunkel G, Nicolasen MJT, Mahata SK, van den Bogaart G. Catestatin as a Target for Treatment of Inflammatory Diseases. Front Immunol. 2018;9:2199.

16. Mahata SK, Corti A. Chromogranin A and its fragments in cardiovascular, immunometabolic, and cancer regulation. Ann N Y Acad Sci. 2019;1455(1):34–58.

17. Muntjewerff EM, Christoffersson G, Mahata SK, van den Bogaart G. Putative regulation of macrophage-mediated inflammation by catestatin. Trends Immunol. 2022;43(1):41–50.

18. Ying W, Mahata S, Bandyopadhyay GK, et al. Catestatin Inhibits Obesity-Induced Macrophage Infiltration and Inflammation in the Liver and Suppresses Hepatic Glucose Production, Leading to Improved Insulin Sensitivity. Diabetes. 2018;67(5):841–848.

19. Kojima M, Ozawa N, Mori Y, et al. Catestatin Prevents Macrophage-Driven Atherosclerosis but Not Arterial Injury-Induced Neointimal Hyperplasia. Thromb Haemost. 2018;118(1):182–194.

20. Mahapatra NR, O’Connor DT, Vaingankar SM, et al. Hypertension from targeted ablation of chromogranin A can be rescued by the human ortholog. J Clin Invest. 2005;115(7):1942–1952.

21. Fung MM, Salem RM, Mehtani P, et al. Direct vasoactive effects of the chromogranin A (CHGA) peptide catestatin in humans in vivo. Clin Exp Hypertens. 2010;32(5):278–287.

22. Biswas N, Gayen J, Mahata M, Su Y, Mahata SK, O’Connor DT. Novel peptide isomer strategy for stable inhibition of catecholamine release: application to hypertension. Hypertension. 2012;60(6):1552–1559.

23. Avolio E, Mahata SK, Mantuano E, et al. Antihypertensive and neuroprotective effects of catestatin in spontaneously hypertensive rats: Interaction with GABAergic transmission in amygdala and brainstem. Neuroscience. 2014;270:48–57.

24. Ying W, Tang K, Avolio E, et al. Immunosuppression of Macrophages Underlies the Cardioprotective Effects of CST (Catestatin). Hypertension. 2021;77(5):1670–1682.

25. Dasgupta A, Bandyopadhyay GK, Ray I, et al. Catestatin improves insulin sensitivity by attenuating endoplasmic reticulum stress: In vivo and in silico validation. Comput Struct Biotechnol J. 2020;18:464–481.

26. Mahata SK, O’Connor DT, Mahata M, et al. Novel autocrine feedback control of catecholamine release. A discrete chromogranin A fragment is a noncompetitive nicotinic cholinergic antagonist. J Clin Invest. 1997;100(6):1623–1633.

27. Mahata SK, Mahapatra NR, Mahata M, et al. Catecholamine secretory vesicle stimulus-transcription coupling in vivo. Demonstration by a novel transgenic promoter/photoprotein reporter and inhibition of secretion and transcription by the chromogranin A fragment catestatin. J Biol Chem. 2003;278:32058–32067.

28. Mahata SK, Mahata M, Fung MM, O’Connor DT. Catestatin: a multifunctional peptide from chromogranin A. Regul Pept. 2010;162(1-3):33–43.

29. Tota B, Angelone T, Mazza R, Cerra MC. The chromogranin A-derived vasostatins: new players in the endocrine heart. Curr Med Chem. 2008;15(14):1444–1451.

30. Theurl M, Schgoer W, Albrecht K, et al. The neuropeptide catestatin acts as a novel angiogenic cytokine via a basic fibroblast growth factor-dependent mechanism. Circ Res. 2010;107(11):1326–1335.

31. Radek KA, Lopez-Garcia B, Hupe M, et al. The neuroendocrine peptide catestatin is a cutaneous antimicrobial and induced in the skin after injury. J Invest Dermatol. 2008;128(6):1525–1534.

32. Hoq MI, Niyonsaba F, Ushio H, Aung G, Okumura K, Ogawa H. Human catestatin enhances migration and proliferation of normal human epidermal keratinocytes. J Dermatol Sci. 2011;64(2):108–118.

33. Stockl V, Blatsios G, Humpel C, et al. Catestatin-like immunoreactivity in the skin and related sensory ganglia. Neuropeptides. 2025;111:102520.

34. Pedri D, Karras P, Landeloos E, Marine JC, Rambow F. Epithelial-to-mesenchymal-like transition events in melanoma. FEBS J. 2022;289(5):1352–1368.

35. Miskolczi Z, Smith MP, Rowling EJ, Ferguson J, Barriuso J, Wellbrock C. Collagen abundance controls melanoma phenotypes through lineage-specific microenvironment sensing. Oncogene. 2018;37(23):3166–3182.

36. Malekan M, Ebrahimzadeh MA, Sheida F. The role of Hypoxia-Inducible Factor-1alpha and its signaling in melanoma. Biomed Pharmacother. 2021;141:111873.

37. Li B, Shen W, Peng H, et al. Fibronectin 1 promotes melanoma proliferation and metastasis by inhibiting apoptosis and regulating EMT. Onco Targets Ther. 2019;12:3207–3221.

38. Tirado-Hurtado I, Fajardo W, Pinto JA. DNA Damage Inducible Transcript 4 Gene: The Switch of the Metabolism as Potential Target in Cancer. Front Oncol. 2018;8:106.

39. Martin A, Salvador F, Moreno-Bueno G, et al. Lysyl oxidase-like 2 represses Notch1 expression in the skin to promote squamous cell carcinoma progression. EMBO J. 2015;34(8):1090–1109.

40. Hutchenreuther J, Vincent KM, Carter DE, Postovit LM, Leask A. CCN2 Expression by Tumor Stroma Is Required for Melanoma Metastasis. J Invest Dermatol. 2015;135(11):2805–2813.

41. Anastas JN, Kulikauskas RM, Tamir T, et al. WNT5A enhances resistance of melanoma cells to targeted BRAF inhibitors. J Clin Invest. 2014;124(7):2877–2890.

42. Shi H, Kong X, Ribas A, Lo RS. Combinatorial treatments that overcome PDGFRbeta-driven resistance of melanoma cells to V600EB-RAF inhibition. Cancer Res. 2011;71(15):5067–5074.

43. Tchurikov NA, Vartanian AA, Klushevskaya ES, et al. Strong Activation of ID1, ID2, and ID3 Genes Is Coupled with the Formation of Vasculogenic Mimicry Phenotype in Melanoma Cells. Int J Mol Sci. 2024;25(17).

44. Sachindra, Larribere L, Novak D, et al. New role of ID3 in melanoma adaptive drug-resistance. Oncotarget. 2017;8(66):110166–110175.

45. Poveda-Garavito N, Orozco Castano CA, Torres-Llanos Y, et al. ID1 and ID3 functions in the modulation of the tumour immune microenvironment in adult patients with B-cell acute lymphoblastic leukaemia. Front Immunol. 2024;15:1473909.

46. Li L, Zhang S, Li H, Chou H. FGFR3 promotes the growth and malignancy of melanoma by influencing EMT and the phosphorylation of ERK, AKT, and EGFR. BMC Cancer. 2019;19(1):963.

47. Talebi A, Dehairs J, Rambow F, et al. Sustained SREBP-1-dependent lipogenesis as a key mediator of resistance to BRAF-targeted therapy. Nat Commun. 2018;9(1):2500.

48. Dong Z, Liu Y, Lu S, et al. Vav3 oncogene is overexpressed and regulates cell growth and androgen receptor activity in human prostate cancer. Mol Endocrinol. 2006;20(10):2315–2325.

49. Chen X, Chen M, Gu X, et al. Roles of KDM5 demethylases in therapeutic resistance of cancers. Epigenetics Chromatin. 2025;18(1):61.

50. Jose A, Shenoy GG, Sunil Rodrigues G, et al. Histone Demethylase KDM5B as a Therapeutic Target for Cancer Therapy. Cancers (Basel). 2020;12(8).

51. Liu X, Zhang SM, McGeary MK, et al. KDM5B Promotes Drug Resistance by Regulating Melanoma-Propagating Cell Subpopulations. Mol Cancer Ther. 2019;18(3):706–717.

52. Weiss MB, Abel EV, Mayberry MM, Basile KJ, Berger AC, Aplin AE. TWIST1 is an ERK1/2 effector that promotes invasion and regulates MMP-1 expression in human melanoma cells. Cancer Res. 2012;72(24):6382–6392.

53. Wu R, Wang C, Li Z, et al. SOX2 promotes resistance of melanoma with PD-L1 high expression to T-cell-mediated cytotoxicity that can be reversed by SAHA. J Immunother Cancer. 2020;8(2).

54. Cheng Q, Wu J, Zhang Y, et al. SOX4 promotes melanoma cell migration and invasion though the activation of the NF-kappaB signaling pathway. Int J Mol Med. 2017;40(2):447–453.

55. Hartman ML, Czyz M. MITF in melanoma: mechanisms behind its expression and activity. Cell Mol Life Sci. 2015;72(7):1249–1260.

56. Vinyals A, Ferreres JR, Campos-Martin R, et al. Regulatory Mechanisms of SPARC Overexpression in Melanoma Progression. Int J Mol Sci. 2025;26(17).

57. Bray F, Laversanne M, Sung H, et al. Global cancer statistics 2022: GLOBOCAN estimates of incidence and mortality worldwide for 36 cancers in 185 countries. CA Cancer J Clin. 2024;74(3):229–263.

58. Boutros A, Croce E, Ferrari M, et al. The treatment of advanced melanoma: Current approaches and new challenges. Crit Rev Oncol Hematol. 2024;196:104276.

59. Miao Y, Quinn TP. Peptide-targeted radionuclide therapy for melanoma. Crit Rev Oncol Hematol. 2008;67(3):213–228.

60. Massaoka MH, Matsuo AL, Figueiredo CR, et al. A novel cell-penetrating peptide derived from WT1 enhances p53 activity, induces cell senescence and displays antimelanoma activity in xeno- and syngeneic systems. FEBS Open Bio. 2014;4:153–161.

61. Soo JK, Castle JT, Bennett DC. Preferential killing of melanoma cells by a p16-related peptide. Biol Open. 2023;12(8).

62. Zhao Z, Li ZQ, Huang YB, et al. An optimized integrin alpha6-targeted peptide capable of delivering toxins for melanoma treatment. J Transl Med. 2025;23(1):495.

63. Yadav V, Zhang X, Liu J, et al. Reactivation of mitogen-activated protein kinase (MAPK) pathway by FGF receptor 3 (FGFR3)/Ras mediates resistance to vemurafenib in human B-RAF V600E mutant melanoma. J Biol Chem. 2012;287(33):28087–28098.

64. DiVito KA, Simbulan-Rosenthal CM, Chen YS, Trabosh VA, Rosenthal DS. Id2, Id3 and Id4 overcome a Smad7-mediated block in tumorigenesis, generating TGF-beta-independent melanoma. Carcinogenesis. 2014;35(4):951–958.

65. Wu S, Naar AM. SREBP1-dependent de novo fatty acid synthesis gene expression is elevated in malignant melanoma and represents a cellular survival trait. Sci Rep. 2019;9(1):10369.

66. Aida S, Sonobe Y, Tanimura H, et al. MITF suppression improves the sensitivity of melanoma cells to a BRAF inhibitor. Cancer Lett. 2017;409:116–124.

67. Bianco M, Gasparri AM, Colombo B, et al. Chromogranin A Is Preferentially Cleaved into Proangiogenic Peptides in the Bone Marrow of Multiple Myeloma Patients. Cancer Res. 2016;76(7):1781–1791.

68. Tatemoto K, Efendic S, Mutt V, Makk G, Feistner GJ, Barchas JD. Pancreastatin, a novel pancreatic peptide that inhibits insulin secretion. Nature. 1986;324(6096):476-478.

69. Aardal S, Helle KB, Elsayed S, Reed RK, Serck-Hanssen G. Vasostatins, comprising the N-terminal domain of chromogranin A, suppress tension in isolated human blood vessel segments. J Neuroendocrinol. 1993;5(4):405–412.

70. Curry WJ, Shaw C, Johnston CF, Thim L, Buchanan KD. Isolation and primary structure of a novel chromogranin A-derived peptide, WE-14, from a human midgut carcinoid tumour. FEBS Lett. 1992;301(3):319–321.

71. Koshimizu H, Cawley NX, Kim T, Yergey AL, Loh YP. Serpinin: A Novel Chromogranin A-Derived, Secreted Peptide Up-Regulates Protease Nexin-1 Expression and Granule Biogenesis in Endocrine Cells. Mol Endocrinol. 2011.

72. Chakraborty S, Karmakar S, Basu M, Kal S, Ghosh MK. The E3 ubiquitin ligase CHIP drives monoubiquitylation-mediated nuclear import of the tumor suppressor PTEN. J Cell Sci. 2023;136(18).

73. Kal S, Chakraborty S, Karmakar S, Ghosh MK. Wnt/beta-catenin signaling and p68 conjointly regulate CHIP in colorectal carcinoma. Biochim Biophys Acta Mol Cell Res. 2022;1869(3):119185.

74. Jati S, Munoz-Mayorga D, Shahabi S, et al. Chromogranin A deficiency attenuates tauopathy by altering epinephrine-alpha-adrenergic receptor signaling in PS19 mice. Nat Commun. 2025;16(1):4703.

