## Supplementary Files for "Catestatin suppresses melanoma progression and drug resistance through multitargeted modulation of signaling pathways"

**Running Title:**

Catestatin Peptide Regulates Melanoma Progression

**Keywords:**

Catestatin; melanoma; peptide therapy; drug resistance, transcriptomics analyses

**Corresponding Authors:**

*Sushil K. Mahata, Ph.D.*
Metabolic Physiology & Ultrastructural Biology Laboratory
Department of Medicine
University of California, San Diego (0732)
9500 Gilman Drive
La Jolla, CA 92093-0732, USA


and

*Satadeepa Kal, Ph.D.*


**Supplementary File 1**

| **Patient ID** | **Sex** | **Tissue type** | **Mutation** | **Treatment regime** | **Metastasis(yes/No-location if known)** |
|---|---|---|---|---|---|
| **K06184** | Male | Brain resection, at diagnosis was found at right calf | BRAF wildtype at diagnosis, c-kit not detected | Ipilimumab, Radiation, Pembrolizumab, Radiation again | Yes (Brain,right thigh) |
| **156681** | Female | Left forearm resection | V600E BRAF | TVEC,Ipilimumab, Decitabine, Vemurafenib. Only the left forearm did not respond. | Yes |
| **128128** | Male | Lymph node resection | Not known | Treatment naive | Yes (lymph node) |

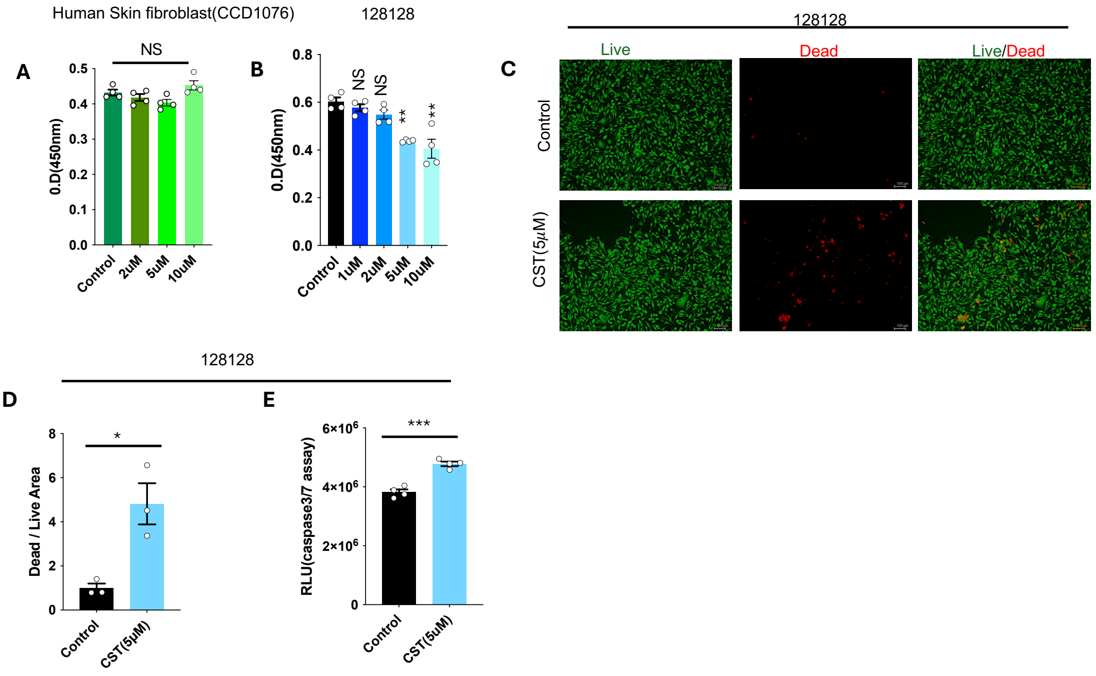


**Supplementary Figure 1**. Anti proliferative effects of CST on melanoma patient derived cell

**(A&B)** Bar graph showing cell viability assay on Human skin fibroblast (CCD1076) and patient derived cell 128128 for 120 hours in response to different concentration of CST (1 μM, 2 μM, 5 μM, and 10 μM) versus vehicle control. (**C)** Micrographs showing green-stained (with calcein AM) live 128128 cells and red-stained (with EthD-III) dead 128128 cells after treatments with CST (5 μM) versus control. Images were captured in Keyence microscope at a magnification of 10X. Scale bar: 100 μm. (**D)** Bar graph showing quantitative analysis of dead/live areas upon CST treatment in 128128 cells. (**E**) Bar graph showing relative light units from caspase 3/7 assay after treatment of 128128 cells with CST or vehicle control. Data were presented as Mean ± SEM and analyzed by Welch's t-test (D&E). **p* ≤ 0.05, ***p* ≤ 0.01, and ****p* ≤ 0.001.

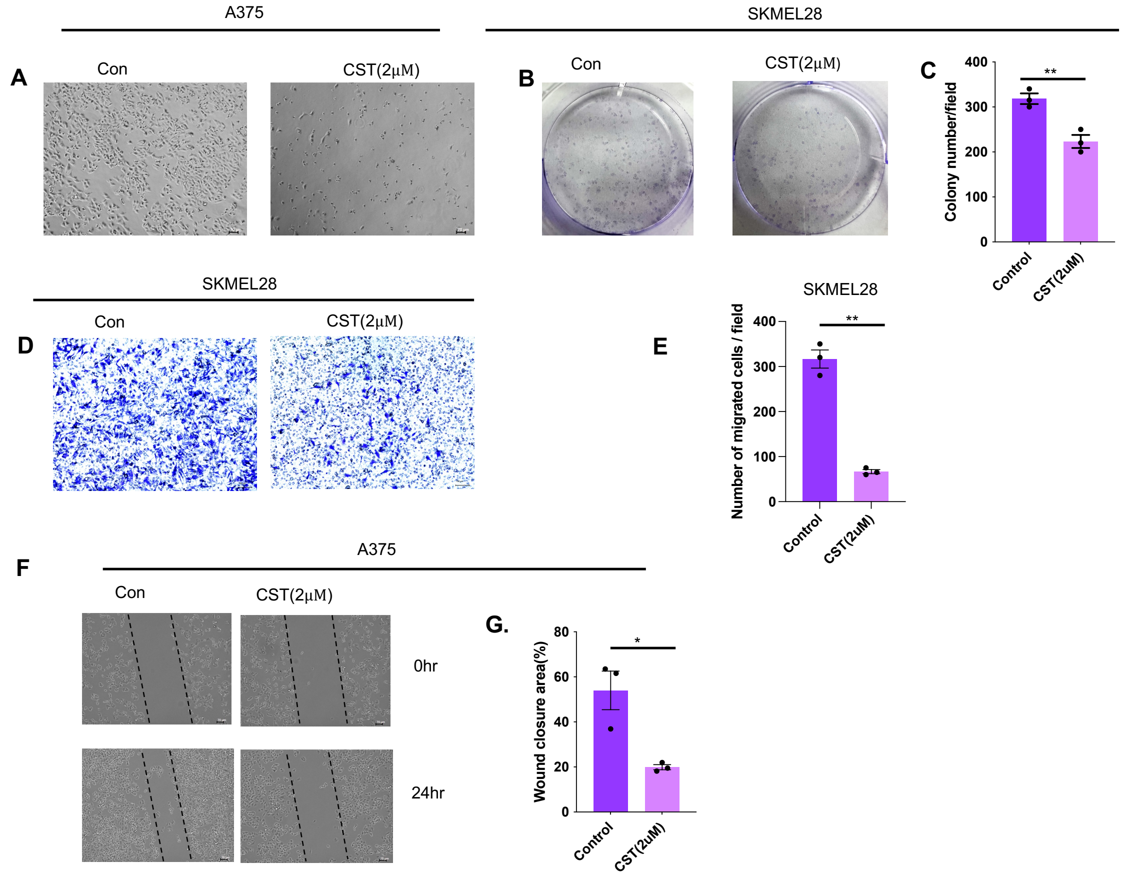


**Supplementary Figure 2**. Melanoma progression inhibition in human melanoma cells upon CST treatment

**(A)** Phase contrast imaging of Control versus CST (2 μM)-treated A375 human melanoma cells for 24 hours. Images were taken at a magnification of 20X. scale bar: 70 μm. **(B&C)** Colony formation assay in SKMEL28 cells after treatments with CST versus control and its quantitative analysis. Brightfield images were captured in of Keyence microscope at a magnification of 10X. Scale bar: 100 μm. (**D&E)** Transwell migration assay in control and CST treated SKMEL28 cells and its quantitative analysis. (**F&G)** Wound healing assay in A375 cells after treatments with CST versus vehicle control for 24 hours and its quantitative analysis. Data were presented as Mean ± SEM and analyzed by Welch's t-test (C, E&G). **p* ≤ 0.05, ***p* ≤ 0.01, and ****p* ≤ 0.001.

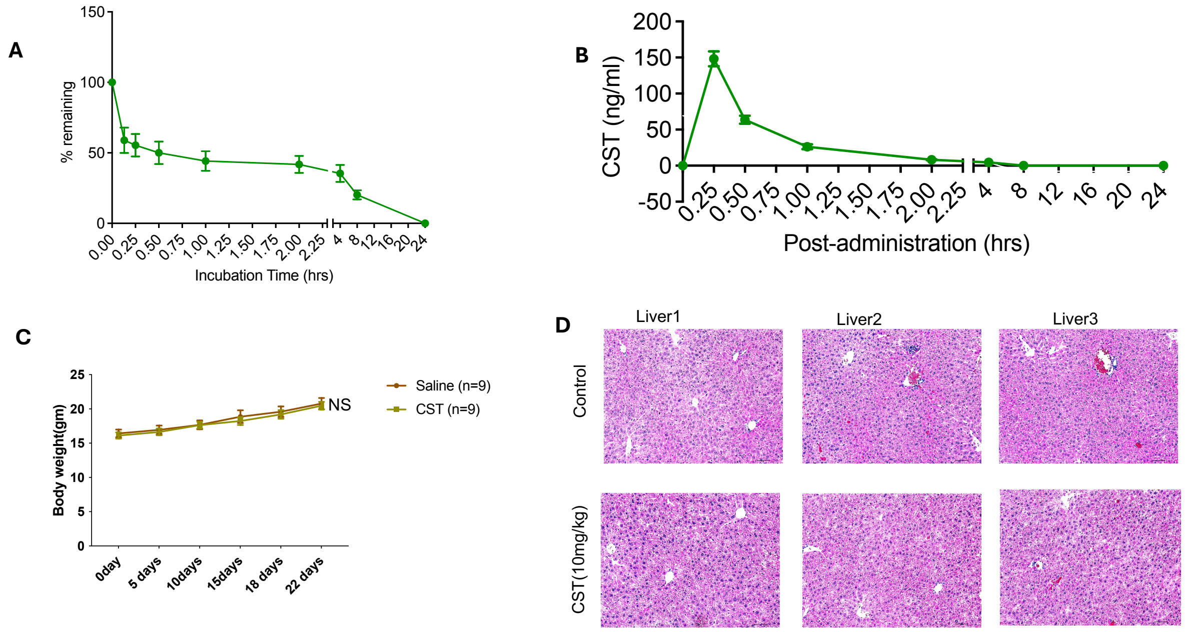


**Supplementary Figure 3.** In vivo melanoma tumor effects upon CST treatment

(**A**) In vitro plasma stability of CST monitored till 24 hours. (**B**) Invivo plasma pharmacokinetics of CST till 24 hours. (**C**) Regulation of body weight by CST (10mg/kg) versus vehicle control in mice bearing B16F10 derived tumors. Data were presented as Mean ± SEM and analyzed by 2-way ANOVA with Sidak's multiple comparison test. **(D)** Histological micrographs of hematoxylin and eosin stained liver sections in mice after treatments with CST versus vehicle control.

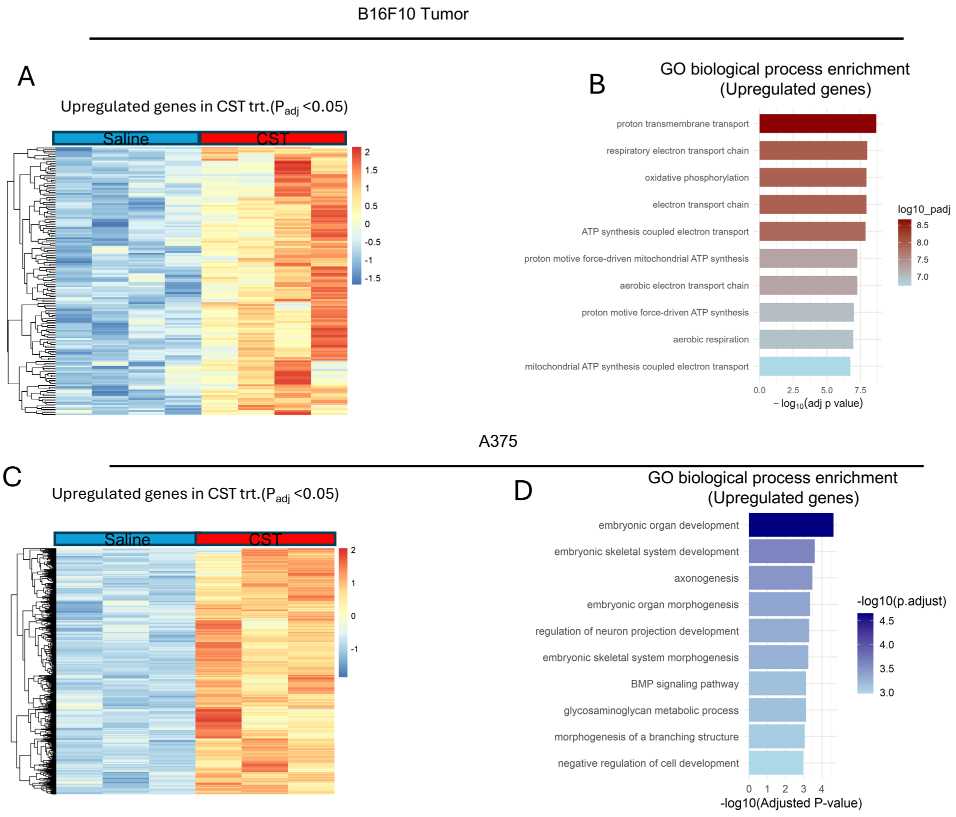


**Supplementary Figure 4.** Molecular mechanism of CST treatment in melanoma tumor and A375 cell line.

**(A)** Heatmap showing upregulated genes upon CST treatment with padj <0.05 in B16F10 tumor (n=4, each group) (**B**) GO analysis showing enriched upregulated pathways in B16F10 tumor in response to CST treatment. (**C**) Heatmap showing upregulated genes in A375 cells after treatments with CST with padj<0.05 (n=3, each group) (**D**) GO analysis showing enriched upregulated pathways in A375 cells upon CST treatment.

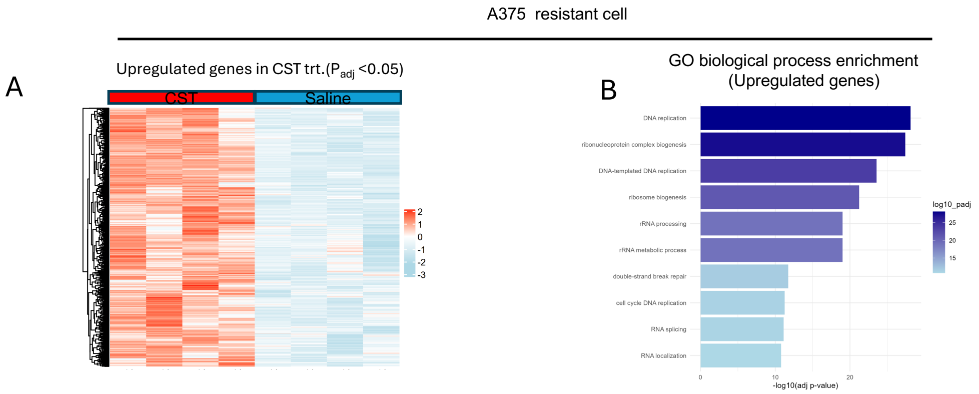


**Supplementary Figure 5.** Molecular mechanism of CST treatment in Vemurafenib resistant melanoma cell A375

**(A)** Heatmap showing upregulated genes in A375 resistant cells upon CST treatment with padj<0.05 (n=4, each group) (**B**) GO analysis showing enriched upregulated pathways in A375 resistant cells in response to CST treatment.
